# Accounting for dispersal improves the understanding of species abundance patterns

**DOI:** 10.1101/2021.03.05.434182

**Authors:** Xiao Feng, Huijie Qiao

## Abstract

A long-standing question in ecology is how are species’ population distributed across space. The highest abundance has been hypothesized to be in the spatial or niche center, though mixed patterns from empirical studies has triggered a recent debate. Here we propose a conceptual framework based on environmental suitability and dispersal to interpret the mixed evidence. We demonstrate that the highest abundance could occur in the spatial center, in the niche center, or somewhere in-between the two centers, depending on the environmental setup and dispersal ability. We found that spatial and niche centers rarely overlap, suggesting the counteracting effect between the two factors, rather than reinforcement, is the norm in determining abundance patterns. The varied locations of highest abundance mirror the mixed evidence in literature, suggesting the “abundant-centre” and “abundant-niche centre” hypotheses are not mutually exclusive. This highlights the importance in understanding the biogeographic patterns through the lens of underlying mechanisms.

## 1. Introduction

### 1.1 The debate of abundance pattern

A long standing question in ecology and biogeography is how are species’ populations distributed across geographic space (Brown 1984). Two hypotheses have been proposed to explain the location with highest abundance. The “abundant-centre” hypothesis predicts that the highest abundance occurs in the center of a species’ geographic range (Hengeveld & Haeck 1982). This hypothesis has received mixed empirical evidence (Tuya *et al.* 2008; Osorio-Olvera *et al.* 2019). The other hypothesis predicts that the abundance is highest in the center of a species’ ecological niche (Maguire 1973; Brown *et al.* 1995; Martínez-Meyer *et al.* 2013; Osorio-Olvera *et al.* 2020). The relationship between niche center and highest abundance is termed “abundant-niche centre” hypothesis, but which also received mixed evidence and is still debated in recent literature (Dallas *et al.* 2017, 2020; Dallas & Hastings 2018; Osorio-Olvera *et al.* 2019; Dallas & Santini 2020; Osorio-Olvera *et al.* 2020).

The mixed empirical evidence for the “abundant-centre” and “abundant-niche centre” hypothesis could be partly explained by differences in data and methodology. Not until recently, few studies have fully included the entire range of a species, dissipating that a species’ geographic range could always be dynamic (Schurr *et al.* 2012), thus limiting the ability to fully test the abundance hypotheses (Fenberg & Rivadeneira 2011; Dallas *et al.* 2017). The rise of citizen-science projects, as well as digitization of museum collections and long-term monitoring networks, notably eBird (Sullivan *et al.* 2009), have greatly improved the data coverage and stimulated a series of interesting investigations (Martínez-Meyer *et al.* 2013; Dallas *et al.* 2017; Osorio-Olvera *et al.* 2020). The differences in methodology among the investigations could be a major explanation for the mixed evidence among the series of explorations, e.g., how the distance is calculated, how the spatial or niche center is quantified, how statistical models are implemented and interpreted (Dallas *et al.* 2017, 2020; Osorio-Olvera *et al.* 2019; Santini *et al.* 2019; Osorio-Olvera *et al.* 2020). Further, the different reflections in the definition, structure, and dimensionality of ecological niche could also lead to different interpretations of the results (Brown 1984; Soberón & Nakamura 2009; Blonder 2016; Soberón & Peterson 2020).

The differences in data and methodology could account for the different conclusions among different studies, but how to explain the mixed evidence from the same study, when the same methodology or workflow was used? A naive answer will be why shouldn’t the observed pattern be case by case, where the observed pattern shall depend on species’ ecological traits (such as dispersal, body size, niche breadth) and the environment (such as habitat heterogeneity, dispersal barrier, and climatic gradient) (Flügge *et al.* 2012; Osorio-Olvera *et al.* 2019; Santini *et al.* 2019; Dallas & Santini 2020; Osorio-Olvera *et al.* 2020). Each aspect of the species and environment can represent a potential underlying mechanism that interacts with other mechanisms and together determine the abundance pattern in the geographic space. Therefore, it is possible that the mechanism underneath the “abundant-centre” and “abundant-niche centre” hypothesis could be both right. However, because multiple underlying mechanisms could interact with each other and together determine the abundance patterns, thus, it becomes difficult to fully prove either hypothesis based on empirical data. In other words, mechanisms could be used to infer empirical patterns, but not vice versa.

### 1.2 Two interacting mechanisms

Following the predecessors’ explorations, we propose an alternative hypothesis to explain the contrary in empirical observations. Our rationale is that the spatial pattern of organisms is analogous to a biological phenotype, which is determined by more than one underlying mechanisms and their interactions (Warren *et al.* 2014), like the classic gene × environment interactions (Relyea & Ricklefs 2013; Lira-Noriega & Manthey 2014). When one mechanism plays a dominant role, the pattern will be more inclined to support the theoretical prediction of this dominating mechanism. When multiple mechanisms counteract each other, the observed pattern will be less favored by either theoretical prediction. Our hypothesis is that environmental suitability and dispersal are two of the major underlying mechanisms that co-determine species’ abundance patterns in geographic space. Therefore, the mixed evidence from empirical studies could represent the interactions between the two mechanisms.

The idea of environmental-suitability-determined abundance has long been proposed and discussed in the literature (Maguire 1973; Brown *et al.* 1995; Martínez-Meyer *et al.* 2013; Osorio-Olvera *et al.* 2020). It assumes the positive association between environmental suitability and population growth rate, and thus the optimum environmental condition (niche center) has the highest abundance (“abundant-niche centre” hypothesis). However, the role of dispersal has received less emphasis in explaining the spatial patterns of abundance in literature (Osorio-Olvera *et al.* 2019; Dallas & Santini 2020). Dispersal is one of the fundamental processes in biogeography, by which biotas respond to spatial and temporal dynamics of geographic template (Lomolino *et al.* 2010). While all other conditions being the same, a species with higher dispersal ability will have higher probability of reaching remote sites and thus potentially a broader geographic distribution. Thus, dispersal is fundamental in determining how biodiversity responses to climate change (Travis *et al.* 2013; Polato *et al.* 2018).

How can dispersal affect the location of the highest abundance? Dispersal, can be thought as a process of spatial redistribution of populations across the landscape (Bohonak 1999; Capinha *et al.* 2015; Fodelianakis *et al.* 2019). For example, in a landscape with uniform environmental suitability where species have uniform population growth, dispersal can affect the species’ abundance pattern through the migrations among different sites. When dispersal is not a limiting factor, each site shall have an equal number of inward and outward migrations, thus dispersal will have no net effect in the abundance pattern. But, when dispersal is a limiting factor that leads to unequal inward and outward migration, the location with highest net inward migrations shall have the highest abundance. In a landscape without dispersal barriers, the spatial center will likely have the highest abundance because of the higher connectivity compared with other locations. Specifically, outward migration will be a fixed quantity pre-determined by dispersal capacity (dispersal function), while inward migration for a focal location will be determined by the number of connections with other locations, which are the source of inward migration. When dispersal is a limiting factor, being in the spatial center means more surrounding locations, as compared to a corner that is partly surrounded by outside unsuitable habitat.

Therefore, in a landscape with more homogeneous environment (e.g., uniform environmental suitability and population growth rate), the spatial center will tend to have the highest abundance when unbalanced migration occurs. On the other hand, when a strong environmental gradient (e.g., higher variance of population growth rates) exists, or when balanced migration occurs, the dispersal-induced redistribution of populations will have less net effect in the abundance pattern. In this case, the observed pattern shall favor the “abundant-niche centre” hypothesis. Finally, in scenarios when neither dispersal nor environmental gradient plays a dominating role, the observed abundance pattern will represent the interactions of the two underlying mechanisms.

### 1.3 Relative position of niche center and spatial center

Still, we have not discussed whether the location of the niche center could overlap the spatial center of a species’ geographic range, which can further affect the observed abundance pattern. In theory, being a niche center does not preclude the possibility of being a spatial center, because the niche center is defined in the environmental space and can be projected between environmental and geographic space; this is also termed the duality of ecological niche (Colwell & Rangel 2009). Generally speaking, we could speculate three scenarios of the geographic location of the niche center: 1) the particular combination of environmental conditions of the niche center may not actually exist in a focal species’ geographic range; 2) the niche center may corresponding to one or multiple locations that are different from spatial center of a focal species’ geographic range; and 3) the niche center may fall in the spatial center of a species’ geographic range. When the two centers overlap, we would expect a more pronounced abundance pattern that supports both “abundant-centre” and “abundant-niche centre” hypothesis; when they do not overlap, we may see an abundance pattern, reflecting the interactions of the two mechanisms.

### 1.4 Overview of objectives

We conduct a series of simulation experiments to demonstrate how the abundance pattern would be affected by environmental suitability and dispersal ability. In particular, we aim to investigate in what scenarios the observed abundance pattern will support the “abundant-centre” or “abundant-niche centre” hypothesis. We also investigate how frequently the location of the niche center overlaps with the spatial center.

## 2. Material and methods

### 2.1 Overview of experiment setup

#### 2.1.1 Virtual landscape

We designed a virtual landscape using a 31-by-31 grid system. The simulated virtual landscape was used to represent a virtual species’ geographic distribution, and the areas outside the focal area were assumed to be unsuitable for the species. Within the virtual landscape, grid cells could have different environmental suitabilities that are positively associated with population growth rate, as commonly assumed by ecological niche theory and “abundant-niche centre” hypothesis (Maguire 1973; Brown *et al.* 1995; Martínez-Meyer *et al.* 2013; Osorio-Olvera *et al.* 2020). Among our simulations, the virtual landscape was assigned varied environmental gradients. The environmental suitability of a location is negatively associated with the spatial distance to the location of niche center (Fig. 1a). This distance-based decay relationship was defined as the strength of the environmental gradient.

**Figure 1.**
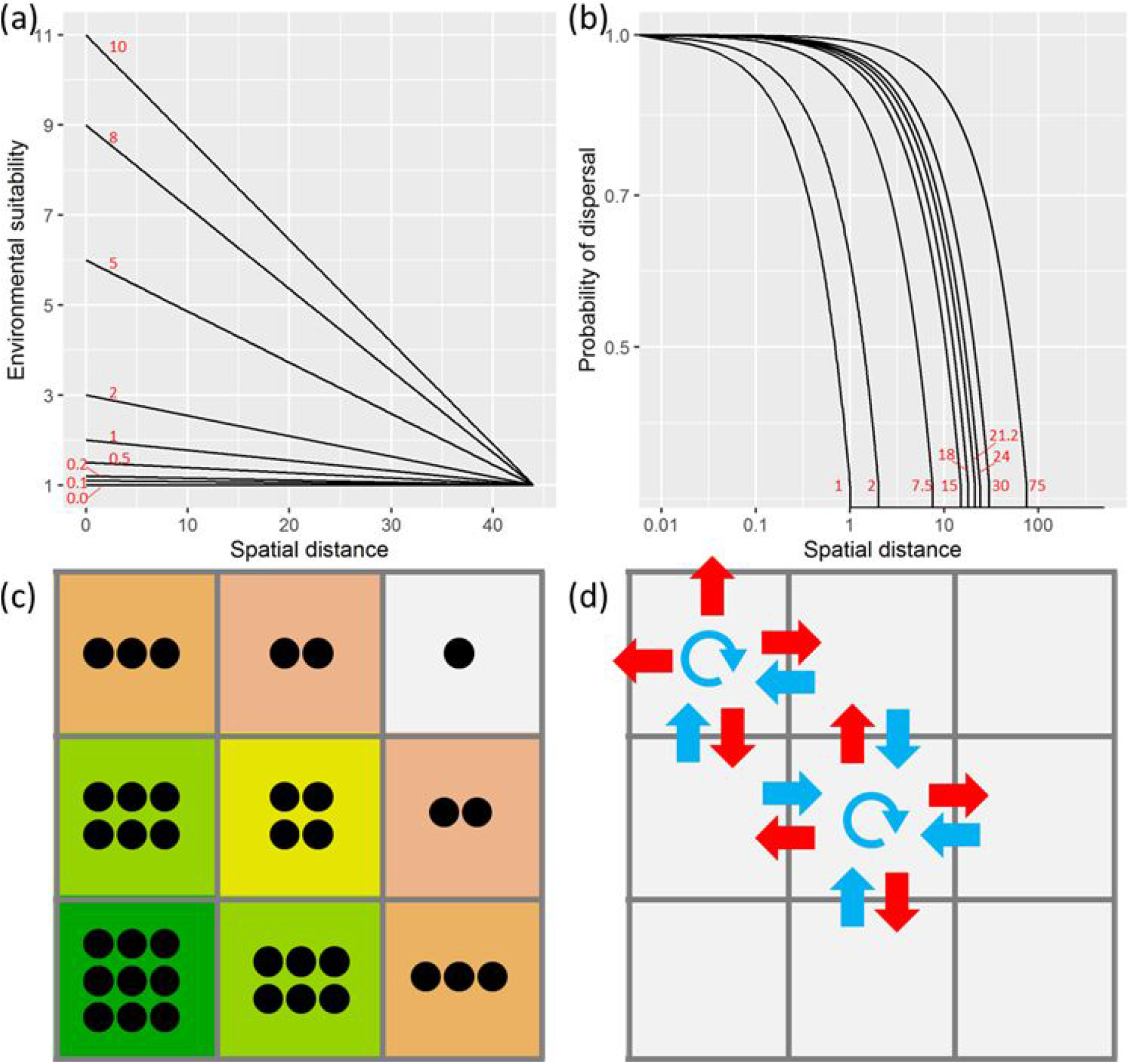
Conceptual illustration of the environmental suitability and dispersal in a virtual landscape. The environmental suitability is highest in the location of the niche center, and linearly decreases over spatial distance (a). The probability of dispersal between two locations decreases exponentially with spatial distance (b). The distance-based decay function is implemented at a gradient of strength for environmental suitability (0-10) and dispersal (1-75). In the virtual landscape, higher environmental suitability is assumed to be associated with higher population growth rate (c). When the dispersal ability is limited (e.g., 1 pixel), the spatial center will have the highest net inward migration (d).

#### 2.1.2 Dispersal

We assume individuals of a species could freely disperse within a maximum distance (D_max_). The probabilities of disperse from cell_i_ to cell_j_, denoted as p_i, j_, decreases exponentially with spatial distance (d_i, j_) following (Osorio-Olvera *et al.* 2019) (Fig. 1b). The p_i, j_ was defined as:

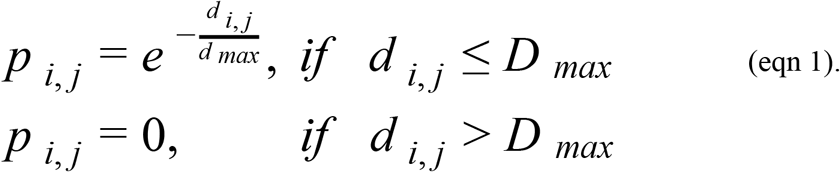

The lowest probability of dispersal occurs when the dispersal distance reaches maximum dispersal distance, beyond which the probability is 0. Individuals are allowed to disperse outside the virtual landscape but will not survive. Therefore, no inward migration will come from cells outside the virtual landscape. In our subsequent experiments, we defined species with different dispersal ability, coupled with varied environmental gradients.

#### 2.1.3 Population growth

We simulated both population growth and dispersal processes in the virtual landscape. The initial population size was set to a constant value (here 961) across all cells. We set the two processes to occur at the same frequency (once per iteration). Within one iteration, we assumed the population growth occurs before dispersal. We implemented 20 iterations for each setup of the environmental suitability and dispersal ability and recorded the mean population status of the last iteration.

In the process of population growth, the population size at cell i (N_i, t_) depends on the population size of the previous iteration (N_i, t-1_) and population growth rate (r_i_), defined as:

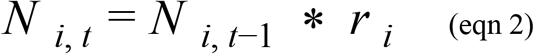

The population growth rate is positively associated with environmental suitability (Fig. 1c). To simplify the simulation we made population growth rate equivalent to the value of environmental suitability, which is ≥ 1. The absolute value of population growth rate or abundance does not correspond to any biological meaning, as the objective of this study is more about the relative abundance pattern (e.g., where is the highest abundance).

During the process of dispersal, the population size at cell_i_ (*N _i, t_*) is calculated as the sum of migration from cells (Δ*N _i,j_*) within the maximum dispersal distance (D_max_), including cell_i_:

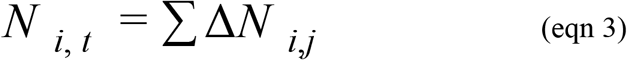

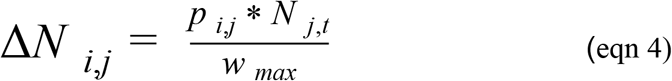

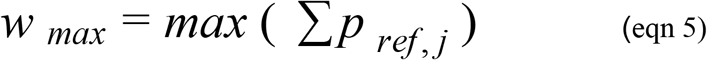

where Δ*Ni*, *j* represents the number of individuals disperse from *cell_j_* to *cell_i_*, and *N_j,t_* represents the number of individuals after the process of population growth at *cell_j_* at iteration *t*. Individuals from cell_i_ will have the highest contribution to the migration to *cell_i_* (analogous to staying at the same cell; Fig. 1d). *W*_*ma*x_ represents a constant weighting factor determined by a defined dispersal ability. *W* is calculated as the sum of dispersal probabilities between a reference cell and all cells surrounding the reference cell within the maximum dispersal ability. *W* reflects the potential of inward migration, and *W_max_* represents the maximum possibility given the environmental conditions and dispersal ability in the virtual landscape. For example, the location of *W_max_* could occur in the spatial center of our virtual landscape when dispersal ability is limited (Fig. 1d).

### 2.2 Simulational experiments

#### 2.2.1 Experiment 1 - one niche center

We conducted a series of experiments with different environmental setups and dispersal abilities. This experiment represents a simplified setup where the environmental suitability decreases linearly from bottom-left corner (location [1,1]) to top-right corner (location [31,31]). The location with highest suitability is used to represent the center of a species’ ecological niche. In this virtual landscape, spatial distance to the location of the niche center has a similar effect as the distance to niche center in environmental space on environmental suitability. In other words, the environmental suitability decreases as either distance increases. The population growth rate at location i (r_i_) is determined by the strength of the environmental gradient (s) and the distance between location i and location [1,1], following this equation:

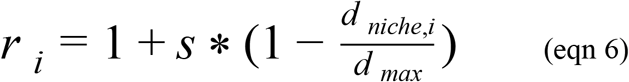

where d_max_ is a constant weighting factor that represents the length of the maximum distance between any two cells in the grid system (i.e., diagonal line in this case).

In this experiment, we designed six levels of the suitability gradients (s = 0, 0.1, 1, 5, 8, 10) and nine levels of dispersal abilities (D_max_=1, 2, 8, 16, 19.2, 22.6, 25.6, 32, 80). A smaller suitability gradient leads to a landscape with more uniform suitabilities, and s = 0 leads to a uniform landscape where all cells have the same suitabilities. The dispersal abilities use the size of the grid system as a reference, for example, 16 represents half of the side length, 32 represents a distance that is just larger than the side length, and 80 represents a dispersal ability that goes beyond the virtual landscape. We conducted (54) simulations based on a full combination of the suitability gradients and dispersal abilities. Each simulation was implemented for 20 iterations, after which we calculated the mean abundance during the last iteration. We recorded the location with the highest abundance and calculated its distance to the niche center in the geographic space.

#### 2.2.2 Experiment 2 - two niche centers

To mimic a slightly more complex landscape, we conducted an additional experiment where multiple (two, in this case) niche centers exist. The niche centers were placed in two corners of landscape, either the bottom left and top right (i.e., diagonal position) or bottom left and bottom right (i.e., same side). In this experiment, we used a similar setup of suitability gradients (s =0.1, 1, 5, 8, 10) and dispersal abilities (D_max_=1, 2, 8, 16, 19.2, 22.6, 25.6, 32, 80). The implementation was the same as Experiment 1.

#### 2.2.3 Experiment 3 - heterogeneous landscape

To mimic a landscape with heterogeneous environmental structures, we conducted an experiment where the environmental suitabilities were randomly selected from a defined range of values (1 and s+1), where s represents the previously defined levels of suitability gradients (s = 0.1, 1, 5, 8, 10). A higher *s* represented a higher contrast between the highest and lowest environmental suitability. The random selection used a uniform distribution and fixed seed.

#### 2.2.4 Experiment 4 - relative position of niche center and spatial center

Previous experiments aimed to demonstrate the interaction between the environmental setup and species’ dispersal ability, when the location of the niche center(s) does not overlap with the spatial center. But, it is unknown whether and to what extent the two centers could overlap with each other in geographic space. Therefore, we conducted an experiment where we randomly selected squared areas from the terrestrial land with varied size, to represent species’ geographic ranges. We evaluated whether the spatial center of an area has environmental conditions similar to the niche center. We also evaluated whether the location of the niche center is close to the spatial center. More details of this simulation are provided in online supporting materials Appendix S1.

To evaluate the overlap status between spatial and niche centers, we quantified the proportion of overlap between spatial center representations and niche center representations (see Appendix S1). The proportion of overlap ranged from 0 (meaning no overlap) to 1 (meaning fully overlap). We conducted ordinary least square regressions using the spatial or environmental distances between spatial center and niche center as the dependent variable and the side length of the square and spatial autocorrelation as independent variables. We calculated Moran’ I to represent the spatial autocorrelation based on 1,000 random samples from a focal square using the first two principal components of 19 bioclim variables. The two independent variables were scaled to zero mean and one standard deviation. When spatial autocorrelation is higher, environmental conditions would be more similar among locations close to each other, thus potentially forming an environmental gradient in geographic space where the spatial center is more likely to be in the middle of the environmental gradient. We hypothesized that the spatial and niche center will be closer (in both spatial and environmental space) when the spatial autocorrelation is stronger. Our analyses were conducted in R (R Development Core Team 2020), using MASS (version 7.3.51.6), ape (version 5.4.1), and raster (version 3.3.13) packages.

## 3 Results

### 3.1 Results of Experiment 1 - one niche center

We found that the location with the highest abundance (Cell_highest_) can fall in the niche center, in the spatial center, or locations in-between the two centers, depending on the combination of environmental gradient of the landscape and the dispersal ability of the virtual species (Figs. 2, 3). We summarized the patterns into different categories: **a)** In a landscape with uniform environmental suitabilities, the spatial center always had the highest abundance regardless of dispersal ability. **b)** In a landscape with weak environmental gradient (s=0.1), the location with the highest abundance (Cell_highest_) is close to the niche center when the dispersal ability is limited (D_max_=1); Cell_highest_ moved away from niche center when the dispersal ability increased, fell in the spatial center when the dispersal ability was close to the half of the diagonal line (D_max_= 16, 19.2, 22.6), and moved back toward the niche center when dispersal ability got larger. **c)** A similar pattern as **(b)** was found with a slightly stronger environmental gradient (s=1), while the difference was that the Cell_highest_ was always in-between niche center and spatial center and never fell on the spatial center. **d**) As the environmental gradient got stronger (S=5,8,10), the locations of the Cell_highest_ were much closer to the niche center, compared with **(c)**, and eventually reached the niche center when the dispersal ability was beyond the size of the virtual landscape (D_max_ = 80).

**Figure 2.**
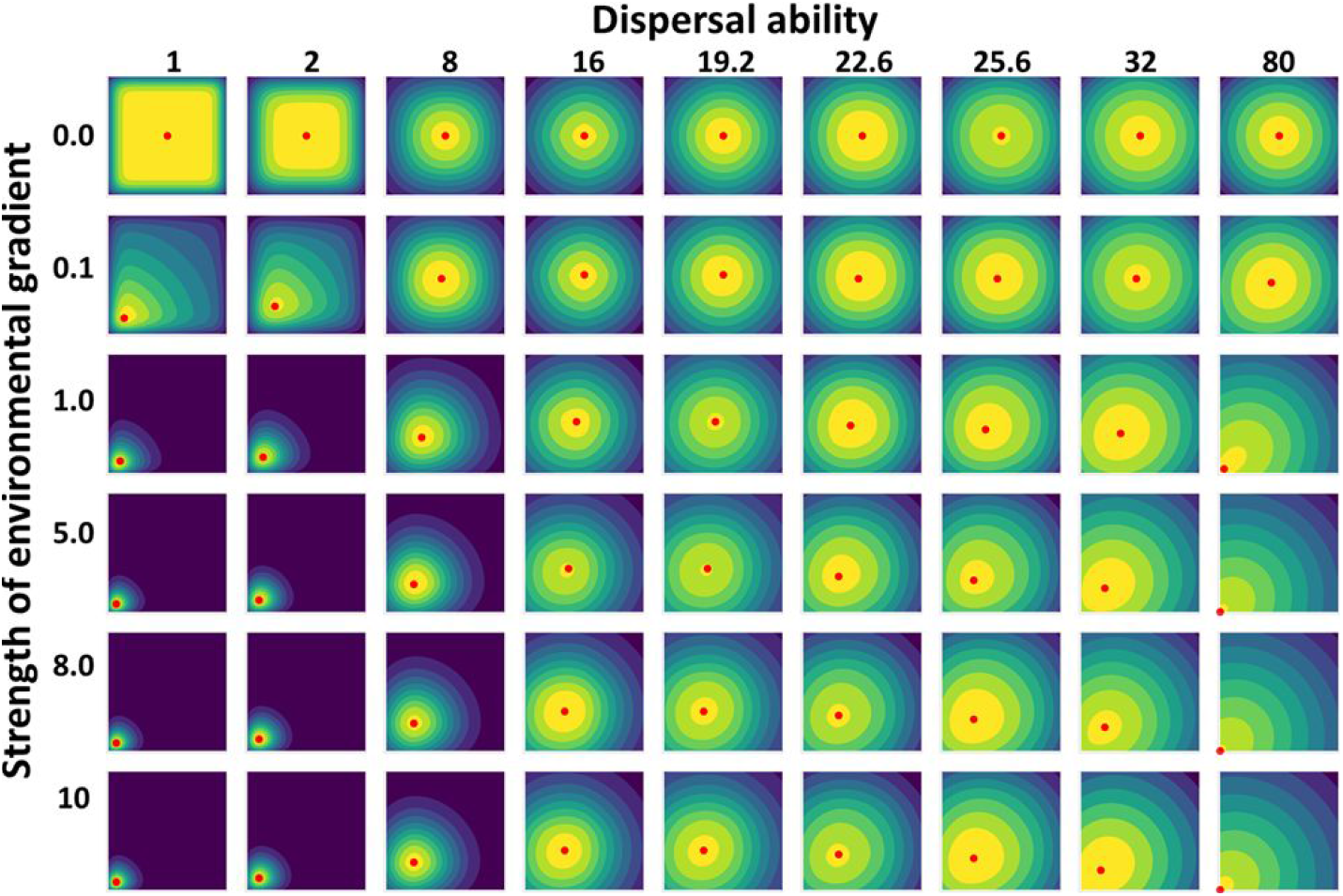
Abundance patterns in a simplified virtual landscape. The panels are arranged by increasing dispersal ability (left to right) and increasing strength of environmental gradient (top to bottom). Within each panel, the abundance values are represented by dark green (low abundance) to yellow (high abundance). The niche center is placed in the bottom-left corner. The red points represent the location with highest abundance.

**Figure 3.**
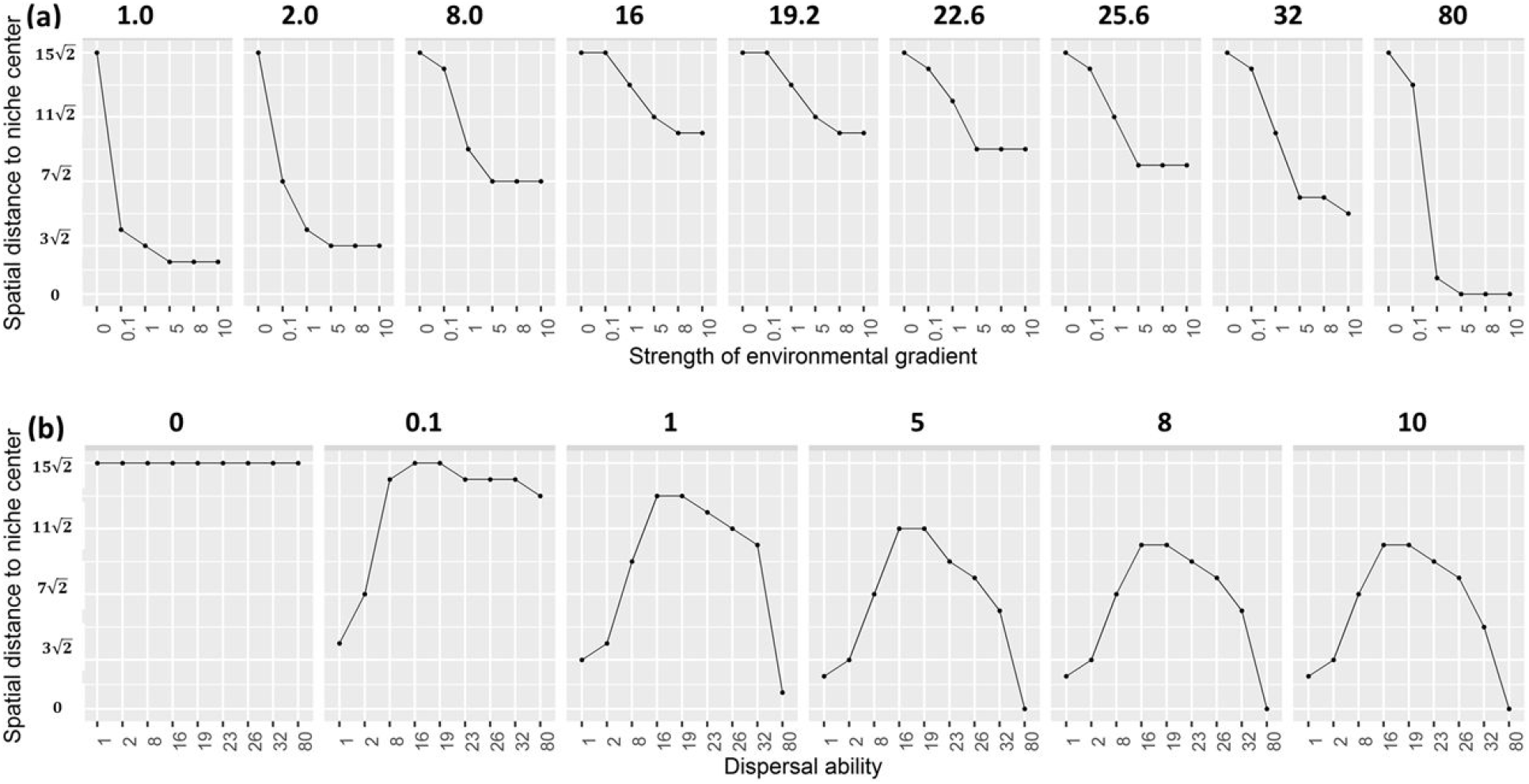
The distances between location of niche center and location with highest abundance in a simplified virtual landscape. Each scatter plot shows how the simulated distance changes (y-axis) along the strength of environmental gradient (a) or dispersal ability (b) on the x-axis. The scatter plots are arranged by different dispersal ability (a; 1-80) or strength of environmental gradient (b; 0-10). The distance (y-axis) also reflects the location of the highest abundance along the diagonal line from bottom-left to top-right: 0 corresponds to location of the niche center [0,0] and 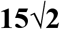 corresponds to spatial center.

To put the results in the context of the debate, the “abundant-niche centre” hypothesis was fully supported when the landscape had a strong gradient of environmental suitability and the species’ dispersal ability was beyond the size of the landscape. When the gradient of environmental suitability existed and the dispersal ability was very weak, the location with highest abundance was close to the niche center, thus partly supporting the “abundant-niche centre” hypothesis. The “abundant-centre” hypothesis was supported in the landscape with no environmental gradient; it can also be supported when the landscape had a weak gradient and the dispersal ability was comparable to the half of the diagonal length of the virtual landscape, a situation the spatial center had relatively higher net inward migration. The results from the simulations well demonstrated that the abundance pattern can be affected by interaction of the two factors. The abundance pattern could favor the prediction of either hypothesis, but could also be different from either prediction. Such mixed patterns found here echo the mixed evidence from empirical studies in the literature.

### 3.2 Results of Experiment 2 - two niche centers

Similar as Experiment 1, we found that the location(s) with the highest abundance (Cell_highest_) can fall in the niche centers, in the spatial center, or locations in-between the two types of centers, depending on the combination of environmental gradient of the landscape and the dispersal ability of the virtual species (Fig. 4a). Compared with Experiment 1, the difference was that two Cell_highest_ can occur at the same time in most of the simulations. In the scenario that the two niche centers were in diagonal position, the two Cell_highest_ lean toward the niche centers when the dispersal ability was week or fell in the niche centers when the dispersal ability was beyond the study area; the two Cell_highest_ lean toward the spatial center when dispersal ability was close to the half of the diagonal line (D_max_= 16, 19.2, 22.6), and the two Cell_highest_ would merged into one when the environmental gradient was week. Compared with the scenario of diagonal position, there were more cases only one Cell_highest_ exist when the niche centers were in the same side of the landscape (Fig. 4b).

**Figure 4.**
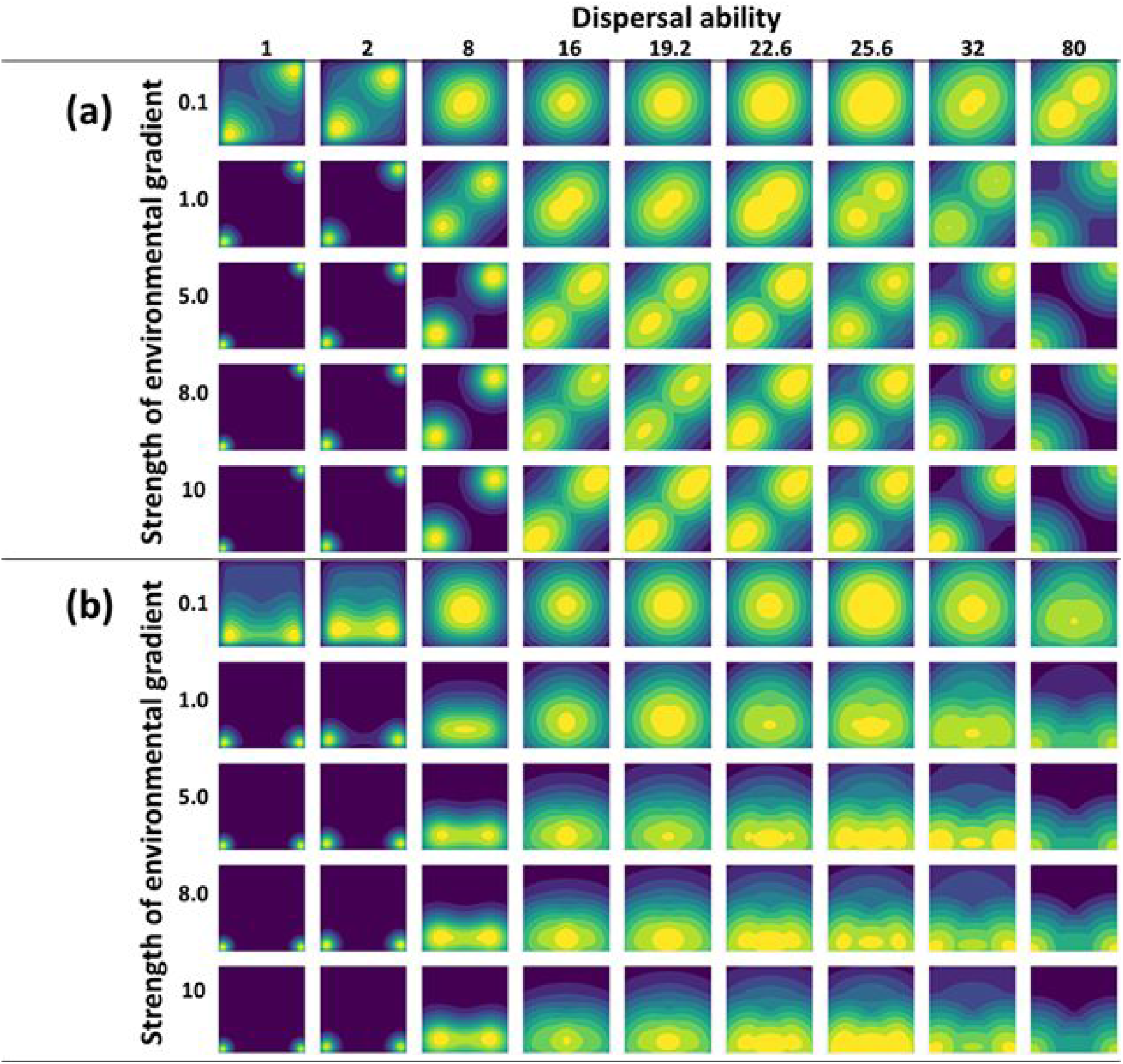
Abundance patterns in a virtual landscape with two niche centers. The panels are arranged by increasing dispersal ability (left to right) and increasing strength of environmental gradient (top to bottom). Within each panel, the abundance values are represented by dark green (low abundance) to yellow (high abundance). The niche centers are placed in the bottom-left and top-right corners (a) or the bottom-left and bottom-right corners (b).

### 3.3 Results of Experiment 3 - heterogeneous landscape

We found that, when dispersal ability was weak (D_max_=1, 2), the locations with high abundances tend to stay in areas with high environmental suitabilities. When dispersal ability was close to the half of the diagonal line (D_max_= 16, 19.2, 22.6), the high abundance locations tend to cluster around the spatial center, and the high abundance locations tend to move away from the spatial center when dispersal ability was beyond the size of the range (Fig. 5).

**Figure 5.**
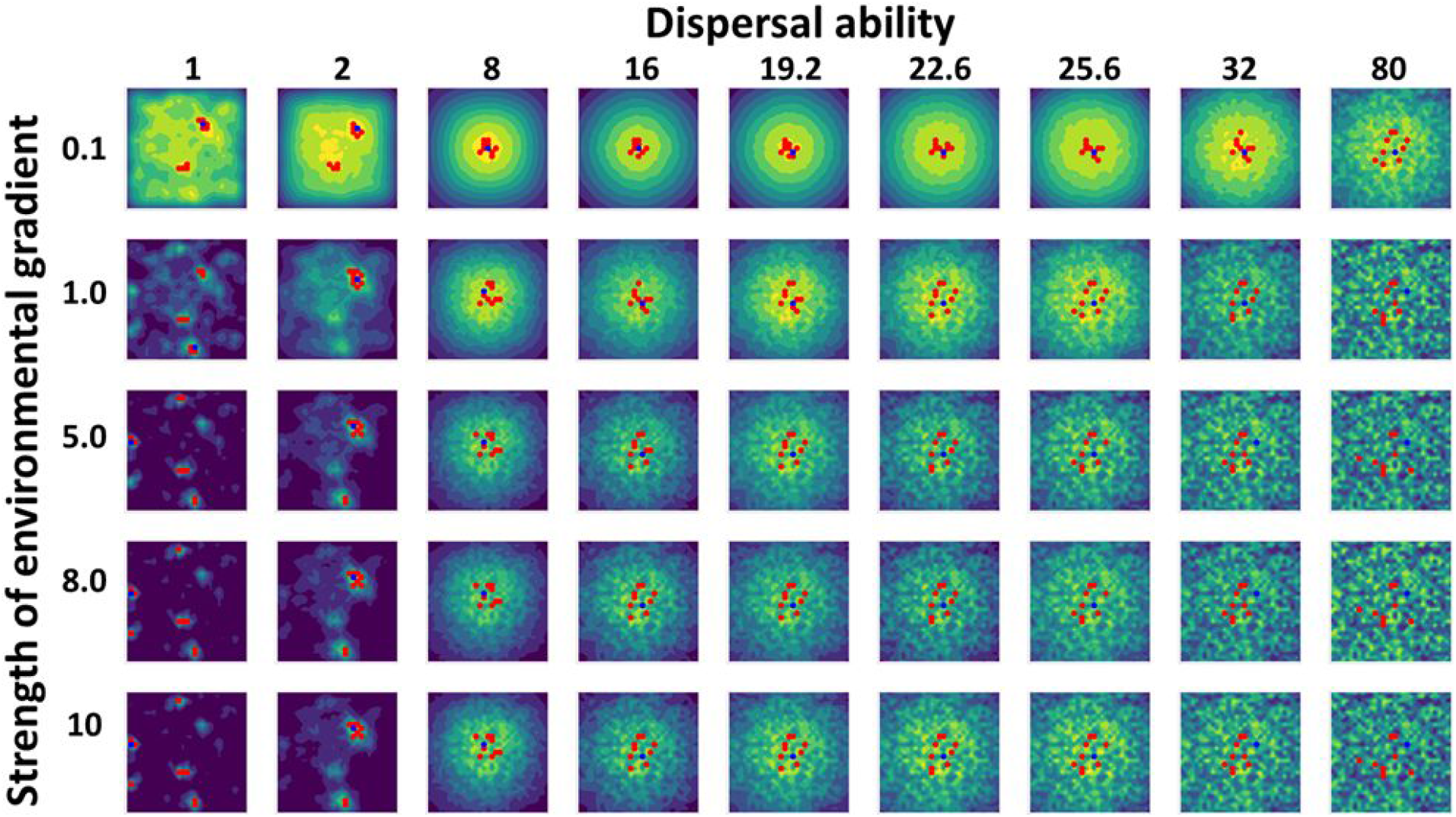
Abundance patterns in a virtual landscape with randomly defined environmental suitabilities. The panels are arranged by increasing dispersal ability (left to right) and increasing strength of environmental gradient (top to bottom). Within each panel, the abundance values are represented by dark green (low abundance) to yellow (high abundance). The blue points represent the location with highest abundance and the red points represent the top ten locations with highest abundance values.

### 3.4 Results of Experiment 4 - relative position of niche center and spatial center

We found that the overlaps between the spatial and niche center representations were mostly low (<10% for 88% of the cases) in both spatial and environmental space, and in rare situations (0.3% of the cases) the overlap could reach 50% (Figs. 6, S1–3). The regressions showed that both spatial and environmental distance between spatial and niche centers were negatively affected by spatial autocorrelation (Moran’s I) and not affected by the size of the square (Table S1).

**Figure 6.**
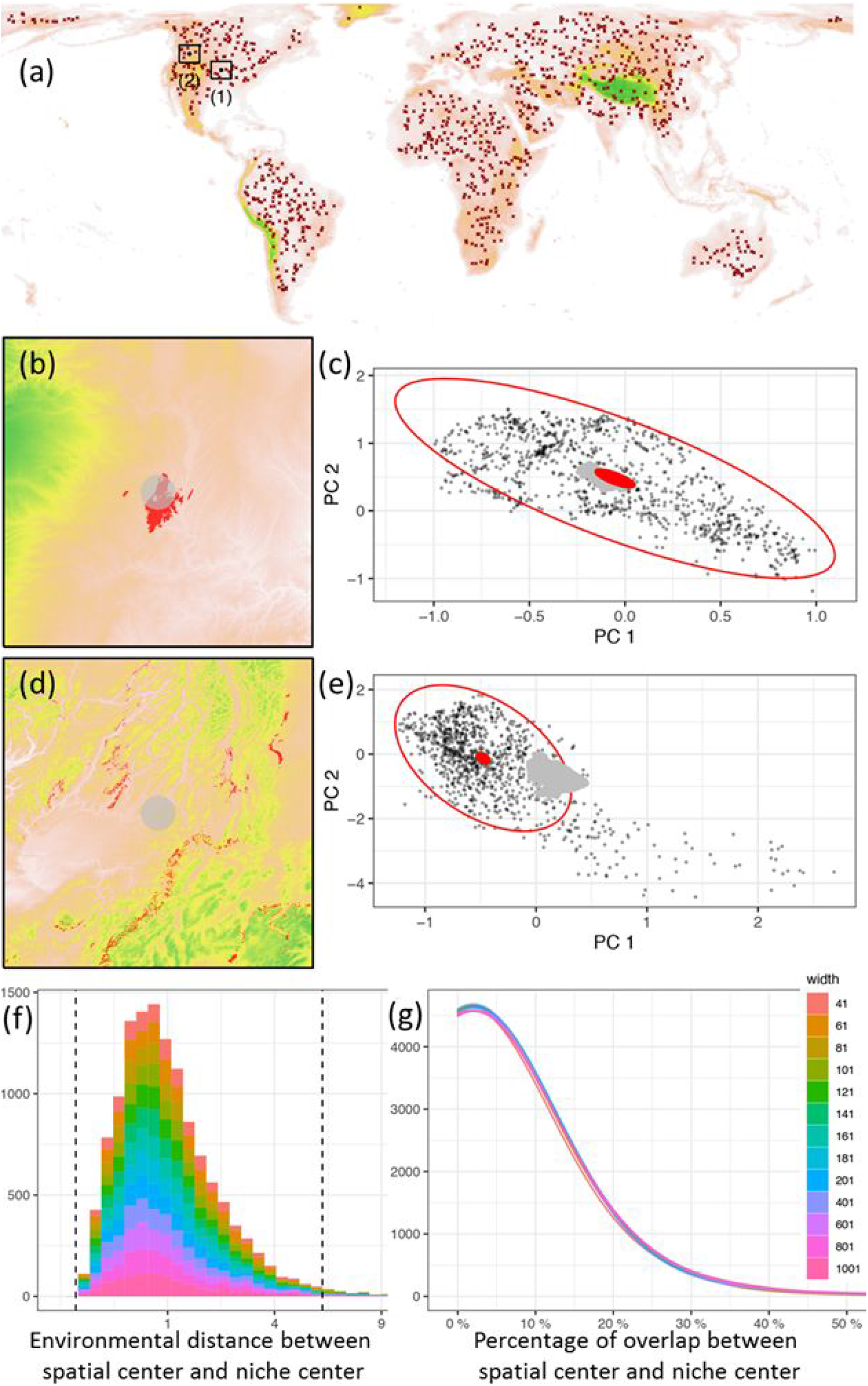
Illustration of the location of niche center and spatial center. Panel a shows the randomly selected initial points (spatial centers) for simulating squares. Two examples of spatial centers are labelled as (1) and (2) in panel a, and are further explained in panels b-e: The spatial center is represented by 1% of points (light gray) surrounding the spatial center in spatial space; the niche center is represented by 1% of points (red) surrounding the niche center in environmental space. The representations of spatial center or niche center are projected to either environmental or spatial space. Among the simulations, some cases (case 1 in panel a) have their spatial center and niche center close to each other in both spatial (b) and environmental space (c), while other cases (case 2) have their spatial center and niche center further away from each other in both spatial (d) and environmental space (e). The gradient of colors in panels a,b,d represent the elevations in the landscape (orange for low elevation and green for high elevation). Panel (f) shows the histogram of distances between spatial center and niche center measured in environmental space. The two dotted lines in panel (f) represent distance of 0.02 and 5.99 to the niche center; the former represents the boundary of the 1% of points surrounding the niche center and the latter represents the boundary of the niche (95% minimum-volume ellipsoid). Panel (g) shows the density plot for the percentage of overlap between the representations of spatial center and niche center.

## 4. Discussion

### 4.1 Understanding the mixed evidence

The spatial center and niche center have been hypothesized to have the highest abundances, though empirical data showed mixed evidences when testing the two hypotheses, thus led to a series of discussions in the recent literature (Dallas *et al.* 2017, 2020; Dallas & Hastings 2018; Osorio-Olvera *et al.* 2019; Osorio-Olvera *et al.* 2020). Our study proposed a conceptual framework to interpret the mixed patterns from empirical studies: dispersal ability and environmental suitability are two of the major drivers that interact with each other and co-determine the spatial distribution of abundances. We demonstrated that the location of the highest abundance could occur in the spatial center, in the location of the niche center, or anywhere in between the two centers, depending on the strength of the dispersal and environmental suitability, and the environmental setup. The varied locations of highest abundance demonstrated here mirrored the mixed abundance patterns from recent studies.

The nonuniform distribution of environmental suitability sets the sense of unequal population growth rate across the geographic space. Dispersal can lead to unequal net inward migration through the process of spatial redistribution of a species’ population. Therefore, it is essentially the environment-determined unequal population growth and dispersal-induced unequal net inward migration, as well as their interactions, that deviate the abundance distribution from uniform and together determine patterns of abundances in a landscape. The spatial center has an advantage of accumulating more net inward migration, because of the overall high connectivity toward areas within a species’ range. The high connectivity is represented by shorter distances between the spatial center and all other locations, while the probability of dispersal is commonly assumed to decay with spatial distance (Nenzén *et al.* 2012; Osorio-Olvera *et al.* 2019). The advantage of more net inward migration is more pronounced for a spatial center when the maximum dispersal distance is close to the radius of a species’ range, and less pronounced when dispersal ability is limited or when the maximum dispersal distance is beyond the diameter of a species’ range. This suggests that the role of dispersal in determining the abundance patterns is relative to the size of a species’ range.

The hypothesized role of dispersal and environmental suitability in determining the abundance pattern allows for several predictions. First, in a landscape with uniform environmental suitability, the locations with highest net inward migration (e.g., the spatial center in the case of no dispersal barriers) will have the highest abundance regardless of dispersal ability. We can also make additional three predictions in a landscape with strong environmental gradient: 1) the location of highest abundance will be close to but not overlap with the location with highest environmental suitability when the dispersal ability is limited; 2) the location with highest environmental suitability will have highest abundance when the maximum dispersal is much larger than the diameter of a species’ range; 3) the location of highest abundance will be furthest away from the location with highest environmental suitability when the maximum dispersal distance is close to the radius of a species’ range. The proposed predictions could be partly supported by the results from (Osorio-Olvera *et al.* 2020). They found that compared with migratory species, the non-migratory species showed stronger negative correlations between abundances and niche-centroid distances with migratory species. If the non-migratory behavior could be interpreted as a weaker ability of spatial redistribution of a population, their results could be interpreted as a relatively weaker dispersal ability that led to a more pronounced signal from environmental suitability. However, limitations in population data across a species’ range and difficulties in the quantification of species dispersal (Lowe & McPeek 2014) could restrict throughout validations of our proposed predictions.

### 4.2 Niche center, spatial center, and spatial autocorrelation

The premise of counteracting effect between dispersal and environmental suitability is that the spatial center does not overlap with the location of the niche center. When they do, the effects of unequal net inward migration and unequal population growth will reinforce each other and likely lead to a stronger abundance gradient. But, how often would the two centers overlap with each other? Our simulation revealed that the spatial or environmental distance between the two centers was negatively associated with spatial autocorrelation. In most cases of the simulations, the two centers do not overlap with each other, suggesting the counteracting effect, rather than reinforcement, between dispersal and environmental suitability, is more likely to be the norm.

### 4.3 The two hypotheses could be both right

The conceptual framework of considering the interaction between dispersal and environmental suitability allows a better understanding of the mixed evidence when testing the “abundant-centre” and “abundant-niche centre” hypothesis. The two hypotheses shall be both supported when the spatial center overlaps with the location of the niche center, though this scenario could be rare. In scenarios where the two centers do not overlap, “abundant-niche centre” hypothesis will be more likely supported when dispersal capacity is limited or goes beyond the general size of a species’ range, and/or when the landscape has a strong environmental gradient that subsequently determines a strong gradient of population growth rate. Similarly, the “abundant-centre” hypothesis will be more likely supported when the maximum dispersal ability is close to the radius of a species’ range, and/or the environmental gradient is weak.

Therefore, the conclusion of hypothesis testing could depend on a few factors. The focal study area (a species’ range) and the environmental composition will determine the position of the niche center relative to the spatial center. The relative size of a species’ geographic range versus the dispersal ability of a species can affect the (unequal) process of the redistribution of a population across the landscape. The relative strength of environmental gradient versus the dispersal ability also affects the relative position of the location with highest abundances.

### 4.4 Limitations

Our simulational experiments were based on a few simplified assumptions. The population growth rate was assumed to be constant for each location, and we did not consider carrying capacity or density-dependent population growth rate (Osorio-Olvera *et al.* 2019). This may lead to overestimation of the effect of environmental suitability on abundance patterns. Also, we used a simple geometry (e.g., square) to represent a species’ geographic where no dispersal barrier exists, and did not consider cases when a species geographic distribution could be composed of multiple discontinuous pieces. Therefore, the geometric center of a species’ range in the real world may not be the location with highest net inward migration. In addition, we assumed that the population growth and dispersal happens at the same frequency, though an increase in frequency of either component will lead to a stronger effect of that component in determining the abundance pattern. Real world species could have varied ratios of the frequencies of the two drivers. A more realistic simulation may refine the relative role of environmental suitability and dispersal. However, the theoretical aspects of our conclusions should still stand.

### 4.5 Final remarks

Our study highlighted the role of dispersal and demonstrated how dispersal can interact with environmental suitability in shaping the spatial distribution of abundances. Our proposed framework can be used to interpret the mixed patterns from empirical studies when testing the “abundant-centre” and “abundant-niche centre” hypotheses. Understanding the role of dispersal and environmental suitability allows us to make predictions of the location with highest abundance, as well as its location relative to the spatial center and niche center. Because the spatial center and the location of the niche center rarely overlap, the counteracting effect between dispersal and environmental suitability, rather than reinforcement, is more likely to be the norm that determines the abundance patterns. Therefore, the location with highest abundance will likely fall in-between the spatial niche and niche center, or in other words, in-between the location(s) with highest inward migration and the location(s) with highest environmental suitability. The “abundant-centre” and “abundant-niche centre” hypotheses are not mutually exclusive and could be both right, though they may not always be supported by empirical data as the spatial distribution of a species can be determined by multiple underlying mechanisms (Lomolino *et al.* 2010). This highlights the importance of resorting to the underlying mechanisms for a better understanding of biogeographic patterns.

---

**Table S1.** Summary of the regression analyses. The distance between spatial center and niche center is the dependent variable, and Moran’s I and side length of a square are independent variables. The geographic distance is standardized by the side length of the square.

| Distance between spatial center and niche center | Intercept | Moran's I | Side length of a square | R<sup>2</sup> | P-value |
| --- | --- | --- | --- | --- | --- |
| Standardized geographic distance | **0.215**<br>**±0.000**<br>*** | **-0.024**<br>**± 0.000**<br>*** | 0.000<br>±0.000 | 0.163 | *** |
| Environmental distance | **1.236**<br>**±0.012**<br>*** | **-0.195**<br>**±0.012**<br>*** | 0.005<br>±0.012 | 0.021 | *** |

\*\*\*p < 0.001

**Figure S1.**
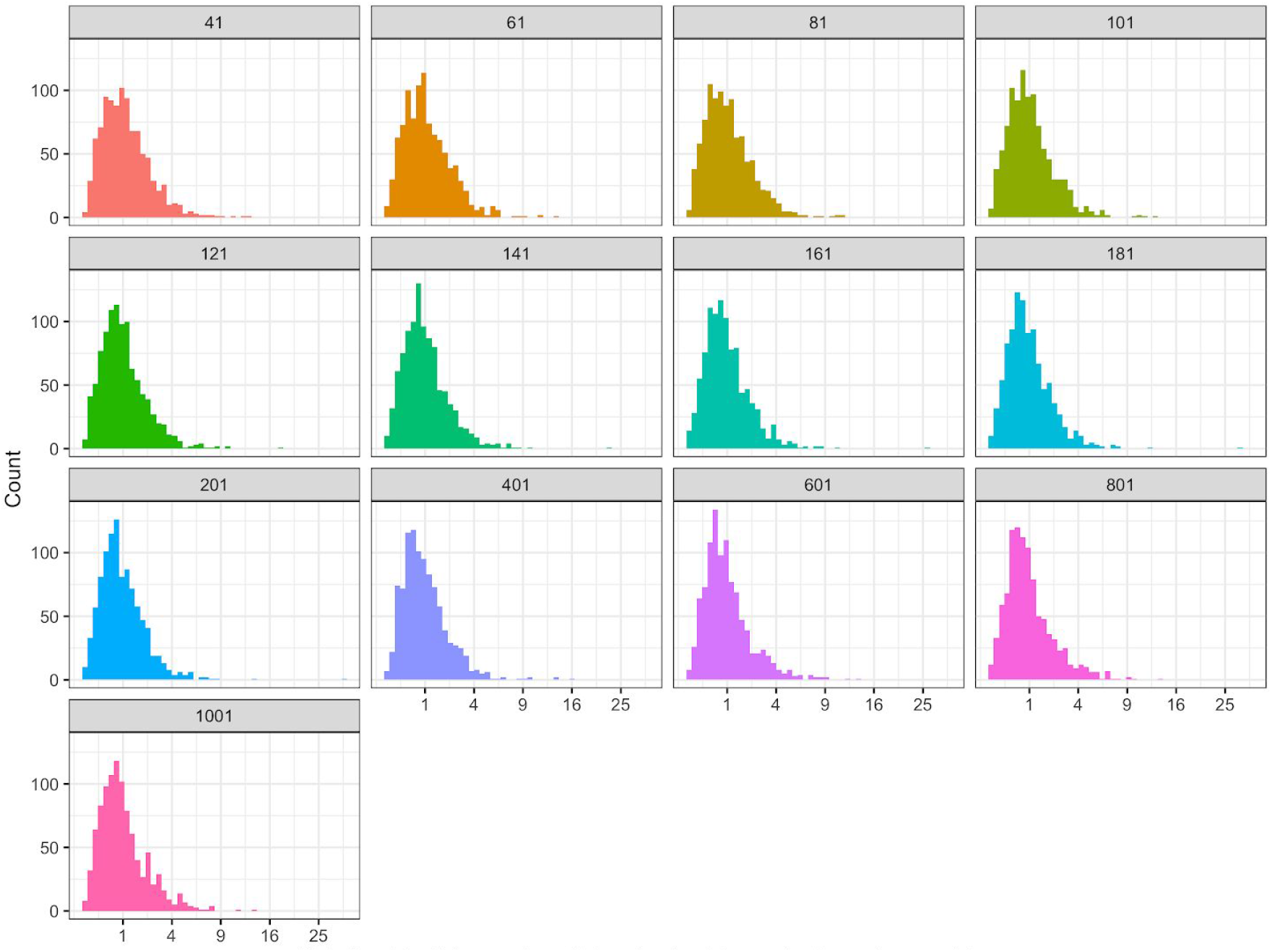
Histogram of environmental distances between spatial center and niche center of randomly selected squares. Different panels represent different side lengths of the squares.

**Figure S2.**
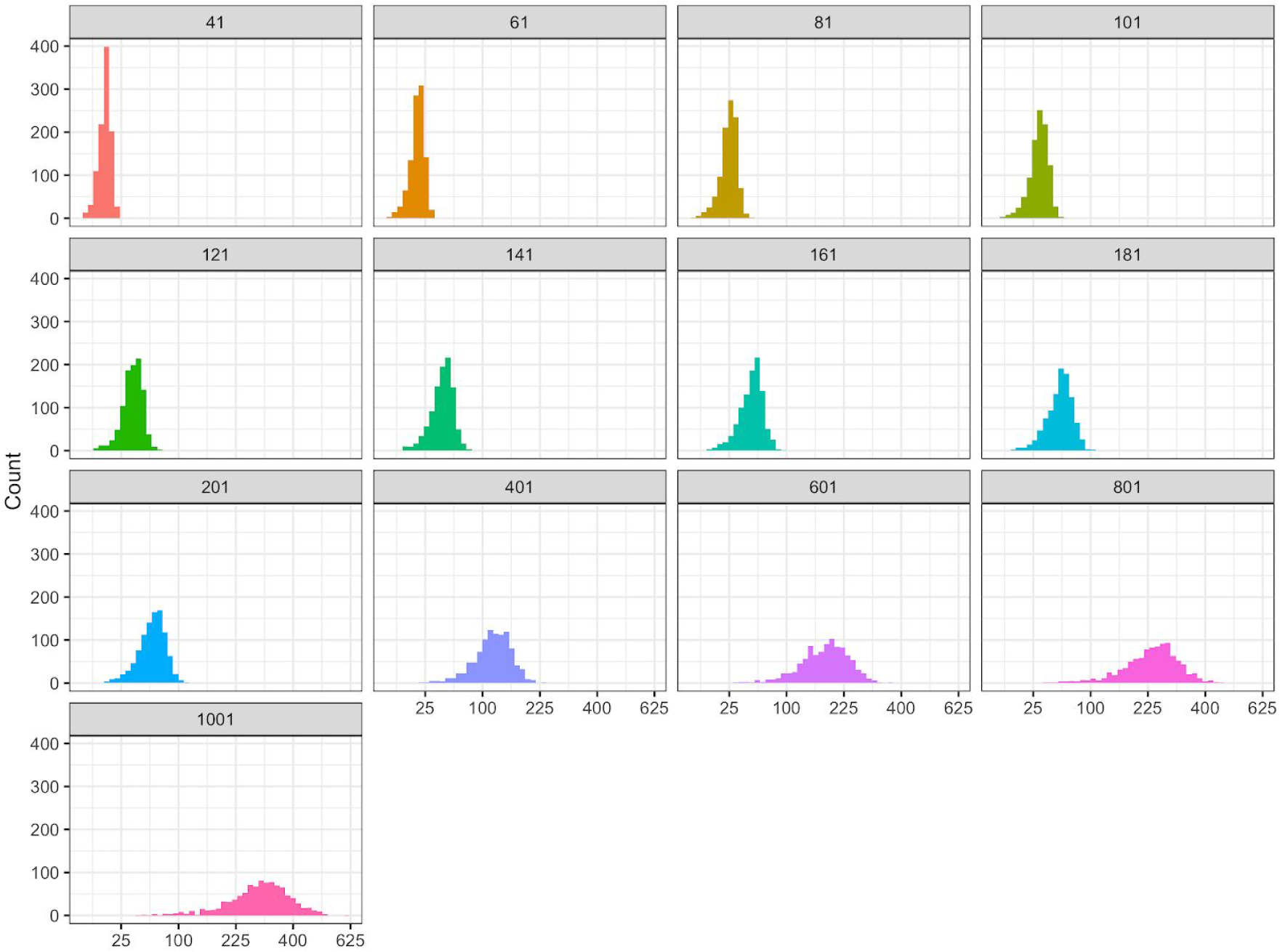
Histogram of spatial distances between spatial center and niche center of randomly selected squares. Different panels represent different side lengths of the squares.

**Figure S3.**
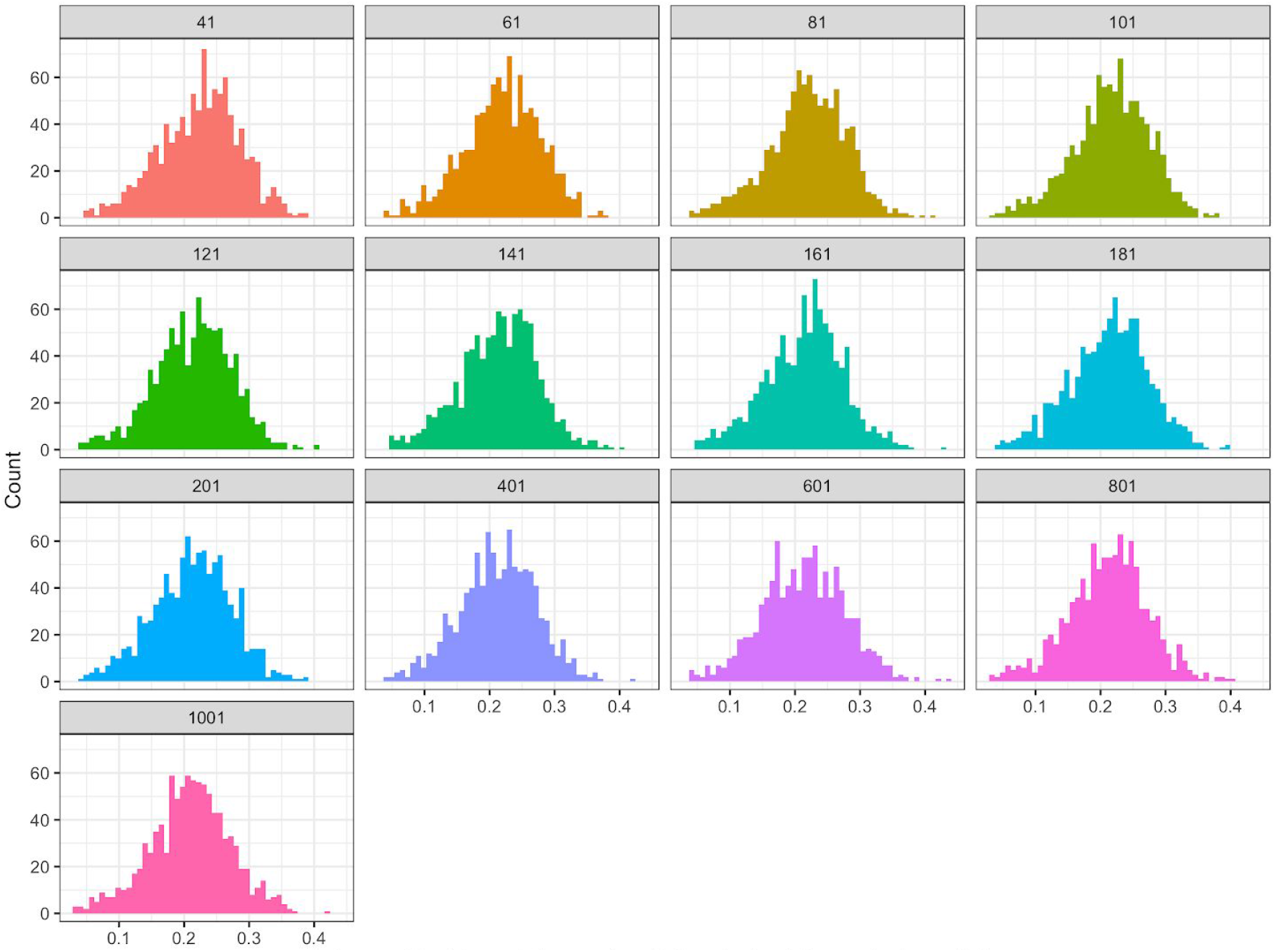
Histogram of standardized spatial distances between spatial center and niche center of randomly selected squares. The spatial distances are divided by the side length of the squares. Different panels represent different side lengths of the squares.

## Appendix S1 Additional details of Experiment 4

Specifically, we first randomly selected a point from the terrestrial land at the resolution of 1km under Eckert IV projection. We used the selected point as the spatial center for a series of squares with different side lengths (41, 61, 81, 101, 121, 141, 161, 181, 201, 401, 601, 801, 1001 km). To avoid the impact of irregular shape on the location of the spatial center (e.g., a square along the coastline), we calculated the proportion of land surface within the 1001 km square and only kept the spatial centers when the proportion of land surface was above 80%. We repeated this process until 1,000 qualified points (spatial centers) were found (Fig. 6a).

Here we considered a square analogous to a species’ geographic distribution. All the pixels within a square were considered as species’ occurrences, based on which we quantified the ecological niche of a focal square using minimum-volume ellipsoid (MVE) (Van Aelst & Rousseeuw 2009; Qiao *et al.* 2016; Osorio-Olvera *et al.* 2020) and the first two principal components of 19 bioclim variables (Fick & Hijmans 2017). The ecological niche of each square was defined as the MVE that covered 95% of the occurrences (Fig. 6b-e).

Instead of identifying the relationship between the spatial and niche center, we resorted to use a group of points surrounding the spatial center in spatial space and environmental conditions surrounding the niche center in environmental space to represent the spatial and niche center. Using spatial or niche center representations could avoid scenarios when the niche center, essentially a combination of environmental conditions, does not exist in spatial space (Soberón & Nakamura 2009), and avoid extreme spatial/environmental distances by using the mean distance. Specifically, the spatial center representations were 1% of the grid cells within the square that were most close to the spatial center; the niche center representations were the 1% of grid cells whose environmental conditions were most close to the niche center in the environmental space. With a larger square, the number of spatial or niche center representations would be larger. The representations of spatial center and niche center were projected to either spatial or environmental space. We calculated the mean spatial distances between the niche center representations and the spatial center in spatial space, and the mean Mahalanobis distances between the spatial center representations and the niche center in environmental space based on the first two principal components of 19 bioclim variables. The mean spatial distances were further standardized by the length of the side length of the square so that the spatial distances were comparable among experiments with different square sizes.

## References

Blonder, B. (2016). Do Hypervolumes Have Holes? Am. Nat., 187, E93–105.

Bohonak, A.J. (1999). Dispersal, gene flow, and population structure. Q. Rev. Biol., 74, 21–45.

Brown, J.H. (1984). On the Relationship between Abundance and Distribution of Species. Am. Nat., 124, 255–279.

Brown, J.H., Mehlman, D.W. & Stevens, G.C. (1995). Spatial Variation in Abundance. Ecology, 76, 2028–2043.

Capinha, C., Essl, F., Seebens, H., Moser, D. & Pereira, H.M. (2015).. The dispersal of alien species redefines biogeography in the Anthropocene. Science, 348, 1248–1251.

Colwell, R.K. & Rangel, T.F. (2009). Hutchinson’s duality: the once and future niche. Proc. Natl. Acad. Sci. U. S. A., 106 Suppl 2, 19651–19658.

Dallas, T.A. & Hastings, A. (2018). Habitat suitability estimated by niche models is largely unrelated to species abundance:Glob. Ecol. Biogeogr., 27, 1448–1456.

Dallas, T.A. & Santini, L. (2020). The influence of stochasticity, landscape structure and species traits on abundant–centre relationships. Ecography, 43, 1341–1351.

Dallas, T., Decker, R.R. & Hastings, A. (2017). Species are not most abundant in the centre of their geographic range or climatic niche. Ecol. Lett., 20, 1526–1533.

Dallas, T., Pironon, S. & Santini, L. (2020). The abundant-centre is not all that abundant: a comment to Osorio-Olvera et al. 2020. bioRxiv.doi:10.1101/2020.02.27.968586

Fenberg, P.B. & Rivadeneira, M.M. (2011). Range limits and geographic patterns of abundance of the rocky intertidal owl limpet, Lottia gigantea. J. Biogeogr., 38, 2286–2298.

Fick, S.E. & Hijmans, R.J. (2017). WorldClim 2: new 1-km spatial resolution climate surfaces for global land areas. Int. J. Climatol., 37, 4302–4315.

Flügge, A.J., Olhede, S.C. & Murrell, D.J. (2012). The memory of spatial patterns: changes in local abundance and aggregation in a tropical forest. Ecology, 93, 1540–1549.

Fodelianakis, S., Lorz, A., Valenzuela-Cuevas, A., Barozzi, A., Booth, J.M. & Daffonchio, D. (2019). Dispersal homogenizes communities via immigration even at low rates in a simplified synthetic bacterial metacommunity. Nat. Commun., 10, 1314.

Hengeveld, R. & Haeck, J. (1982). The Distribution of Abundance. I. Measurements. J. Biogeogr., 9, 303–316.

Lira-Noriega, A. & Manthey, J.D. (2014). Relationship of genetic diversity and niche centrality: a survey and analysis. Evolution, 68, 1082–1093.

Lomolino, M.V., Riddle, B.R., Whittaker, R.J. & Brown, J.H. (2010). Biogeography. Sinauer Associates.

Lowe, W.H. & McPeek, M.A. (2014). Is dispersal neutral? Trends Ecol. Evol., 29, 444–450.

Maguire, B. (1973). Niche Response Structure and the Analytical Potentials of Its Relationship to the Habitat. Am. Nat., 107, 213–246.

Martínez-Meyer, E., Díaz-Porras, D., Peterson, A.T. & Yáñez-Arenas, C. (2013). Ecological niche structure and rangewide abundance patterns of species. Biol. Lett., 9, 20120637.

Nenzén, H.K., Swab, R.M., Keith, D.A. & Araújo, M.B. (2012). demoniche - an R-package for simulating spatially-explicit population dynamics. Ecography, 35, 577–580.

Osorio-Olvera, L., Lira-Noriega, A., Soberón, J., Peterson, A.T., Falconi, M., Contreras-Díaz, R.G., et al. (2020). ntbox : An r package with graphical user interface for modelling and evaluating multidimensional ecological niches. Methods Ecol. Evol., 11, 1199–1206.

Osorio-Olvera, L., Soberón, J. & Falconi, M. (2019). On population abundance and niche structure. Ecography, 42, 1415–1425.

Osorio-Olvera, L., Yañez-Arenas, C., Martínez-Meyer, E. & Peterson, A.T. (2020). Relationships between population densities and niche-centroid distances in North American birds. Ecol. Lett., 23, 555–564.

Polato, N.R., Gill, B.A., Shah, A.A., Gray, M.M., Casner, K.L., Barthelet, A., et al. (2018). Narrow thermal tolerance and low dispersal drive higher speciation in tropical mountains. Proc. Natl. Acad. Sci. U. S. A., 115, 12471–12476.

Qiao, H., Peterson, A.T., Campbell, L.P., Soberón, J., Ji, L. & Escobar, L.E. (2016). NicheA: creating virtual species and ecological niches in multivariate environmental scenarios. Ecography, 39, 805–813.

R Development Core Team. (2020). R: a language and environment for statistical computing.

Relyea, R. & Ricklefs, R.E. (2013). The Economy of Nature: Seventh Edition. Macmillan Learning.

Santini, L., Pironon, S., Maiorano, L. & Thuiller, W. (2019). Addressing common pitfalls does not provide more support to geographical and ecological abundant-centre hypotheses. Ecography, 42, 696–705.

Schurr, F.M., Pagel, J., Cabral, J.S., Groeneveld, J., Bykova, O., O’Hara, R.B., et al. (2012). How to understand species’ niches and range dynamics: a demographic research agenda for biogeography. J. Biogeogr., 39, 2146–2162.

Soberón, J. & Nakamura, M. (2009). Niches and distributional areas: concepts, methods, and assumptions. Proc. Natl. Acad. Sci. U. S. A., 106 Suppl 2, 19644–19650.

Soberón, J. & Peterson, A.T. (2020). What is the shape of the fundamental Grinnellian niche? Theor. Ecol., 13, 105–115.

Sullivan, B.L., Wood, C.L., Iliff, M.J., Bonney, R.E., Fink, D. & Kelling, S. (2009). eBird: A citizen-based bird observation network in the biological sciences. Biol. Conserv., 142, 2282–2292.

Travis, J.M.J., Delgado, M., Bocedi, G., Baguette, M., Bartoń, K., Bonte, D., et al. (2013). Dispersal and species’ responses to climate change. Oikos, 122, 1532–1540.

Tuya, F., Wernberg, T. & Thomsen, M.S. (2008). Testing the “abundant centre” hypothesis on endemic reef fishes in south-western Australia. Mar. Ecol. Prog. Ser., 372, 225–230.

Van Aelst, S. & Rousseeuw, P. (2009). Minimum volume ellipsoid: Minimum volume ellipsoid. Wiley Interdiscip. Rev. Comput. Stat., 1, 71–82.

Warren, D.L., Cardillo, M., Rosauer, D.F. & Bolnick, D.I. (2014). Mistaking geography for biology: inferring processes from species distributions. Trends Ecol. Evol., 29, 572–580.

